# The accuracy of protein structures in solution determined by AlphaFold and NMR

**DOI:** 10.1101/2022.01.18.476751

**Authors:** Nicholas J. Fowler, Mike P. Williamson

## Abstract

In the recent CASP (Critical Assessment of Structure Prediction) competition, AlphaFold2 performed outstandingly. Its worst predictions were for NMR structures, which has two alternative explanations: either the NMR structures were poor, implying that AlphaFold may be more accurate than NMR; or there is a genuine difference between crystal and solution structures. Here, we use the program ANSURR, which measures the accuracy of solution structures, and show that one of the NMR structures was indeed poor. We then compare AlphaFold predictions to NMR structures, and show that AlphaFold tends to be more accurate than NMR ensembles, in particular correctly more rigid in loops. There are however some cases where the NMR ensembles are more accurate. These tend to be dynamic structures where AlphaFold had low confidence. We suggest that AlphaFold could be used as the model for NMR structure refinements, and that AlphaFold structures validated by ANSURR require no further refinement.

## Introduction

In November 2020 the results of the 14^th^ Critical Assessment of Structure Prediction competition (CASP14) revealed that AlphaFold2 (AF2), an AI developed by DeepMind^1^, performed significantly better than all other methods^2,3^. Impressively, the majority of predictions obtained a GDT_TS (Global Distance Test Total Score) score above 80, with a median value of 92.4, where perfect agreement would be 100. Only 5 of the 93 AF2 predictions had a GDT_TS score below 70. Three of these were chains from complexes and two were solved using nuclear magnetic resonance (NMR). Reduced performance for the former was to be expected as AF2 was not designed to predict structural changes that occur from complex formation. Why AF2 did less well for the NMR structures is less obvious. Most NMR structures are small single-chain proteins - a type of structure that should be relatively easy to predict. A possible explanation is that NMR structures are generally of poor quality, implying that AF2 predictions may be more reliable than NMR structures. However, a diametrically opposite explanation is that AF2 is less reliable for predicting NMR structures because it was trained using crystal structures, the assumption being that NMR structures are different from crystal structures because they are obtained in solution at close to body temperature, not in a crystal and (usually) at low temperature^4^.

This raises several important questions: How good is AF2 at predicting <u>solution</u> structures? Is it worth trying to determine NMR solution structures if AF2 structures are as good or better? Are solution structures genuinely different from crystal or AF2 structures? Are NMR structures of good enough quality and reliability to be used as models for the “true” solution structure, and if so, how? This paper aims to provide answers to these questions.

A fundamental problem dating back to the first NMR protein structure^5^ is that there is no reliable way to tell if an NMR structure is correct, ie close to the “true” solution average. The *de facto* method for validating an NMR structure is to compare it to a crystal structure. Surveys carried out based on such comparisons have shown that NMR structures are similar to crystal structures, but in general less well defined (less precise) and also less accurate^6,7^. However, if there are genuine differences between crystal structures and solution structures (for example due to increased flexibility in solution and at higher temperatures), then such comparisons will be misleading. We recently developed a method ANSURR, Accuracy of NMR Structures Using RCI and Rigidity, which calculates the local rigidity of a protein structure^8^, and compares it to the local rigidity as measured using a version of the Random Coil Index^9^ based on backbone NMR chemical shifts^10,11^. The method has been tested on a wide range of structures and provides a reliable guide to accuracy. We have therefore applied ANSURR to answer the questions posed above.

The paper is structured as follows. Firstly, we compare the accuracy of three NMR targets and the corresponding predicted structures from the CASP14 competition, with consideration of both global and local aspects of accuracy. Next, we expand our study to compare 904 structures of human proteins from the AlphaFold Protein Structure Database^12^ with NMR structures from the Protein Data Bank (PDB), highlighting instances where NMR structures are significantly more accurate than AF2 models and *vice versa*. Finally, we investigate the relationship between the estimated accuracy of AF2 models (as predicted by AF2 alongside a structure) with the accuracy determined by ANSURR.

## Results

### The accuracy of target NMR structures and predicted structures from CASP14

ANSURR works by computing two measures of protein flexibility; one obtained from backbone chemical shifts and the other from a structure using the mathematical theory of rigidity. The two measures are compared by computing the rank Spearman correlation coefficient and root-mean-squared deviation (RMSD) between them. The percentile of each value relative to those for all NMR structures in the PDB is used to obtain two scores, termed correlation score and RMSD score, respectively. These scores can be visualised on a single plot so that the best scoring structures (with good correlation and RMSD scores) appear in the top right-hand corner of the plot and the worst scoring appear in the lower left-hand corner (with poor correlation and RMSD scores). CASP14 had three NMR ensembles that were used as targets. These are shown in Figure 1, using either all the structures in the predicted or experimental ensemble (Fig 1a), or the scores averaged across all members of the ensemble (Fig 1b). ANSURR scores for all NMR and AF2 models are provided in supplementary information. One of these (T1055) had AF2 CASP14 predictions that were close to the NMR target structures. However, the other AF2 predictions were very different, with one being worse than the NMR target (T1027), and one being significantly better (T1029). These two structures are now examined in more detail.

**Figure 1.**
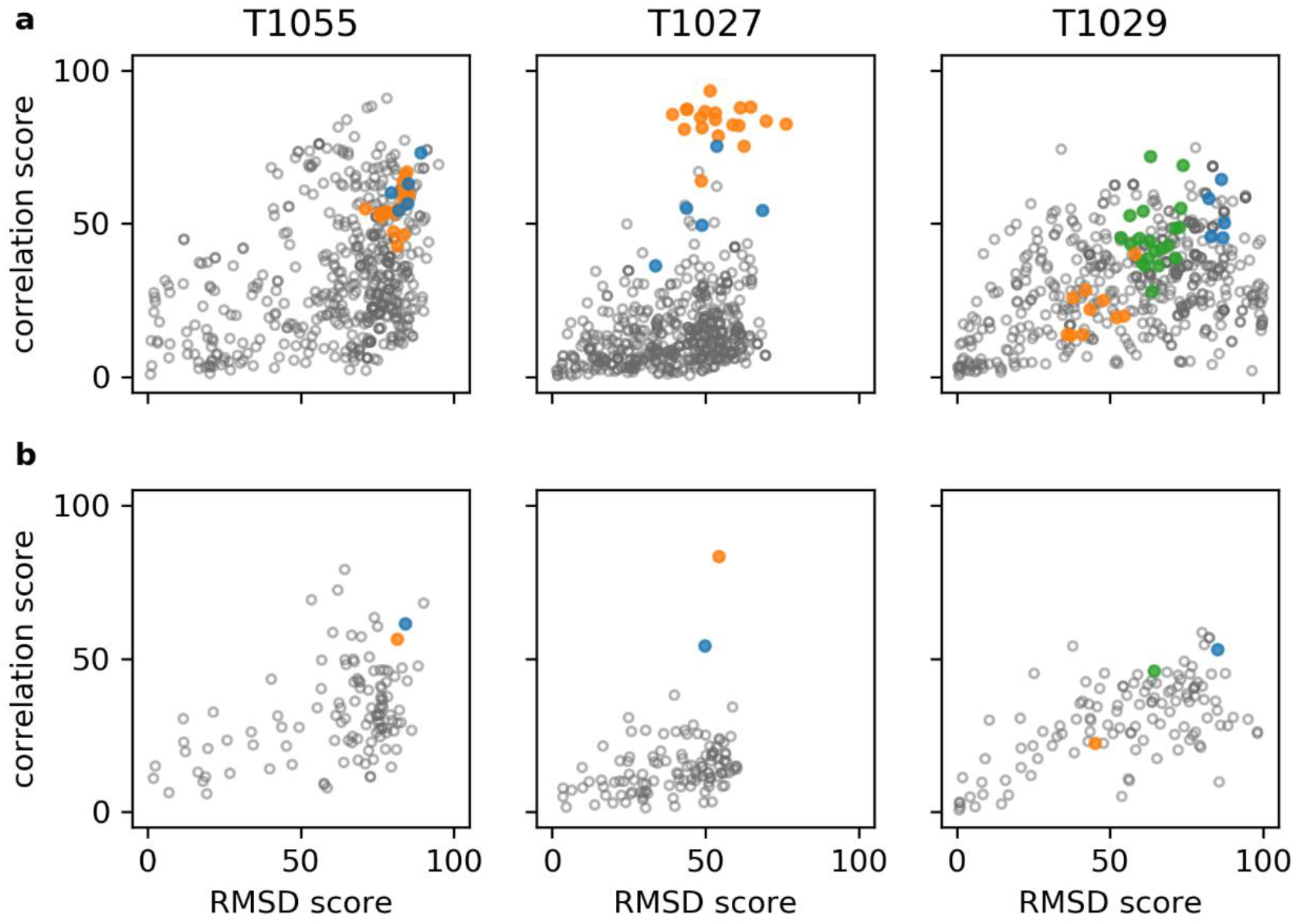
ANSURR scores for the three CASP14 NMR targets. Results for (a) all models and (b) ensemble averages are shown. NMR structures are in orange, AF2 models in blue, and all other predictions in grey. The green points shown for T1029 are scores for an NMR ensemble that was recalculated after the CASP14 results were released, and are discussed below. The NMR structure for T1055 (PDB 6zyc) has 20 models and the NMR structure for T1027 (PDB 7d2o) has 19 models. The original NMR structure for T1029 (PDB 6uf2) has 10 models and the recalculated structure (PDB 7n82) has 20 models. Each group competing in CASP14 could provide up to 5 predictions.

### Target T1027

For target T1027, the target NMR ensemble is more accurate than all predicted structures. However, the AF2 models are the best scoring of the predicted structures, with one model approaching the accuracy of the NMR ensemble. Thus far, it is a fairly unremarkable result. However, interesting lessons can be learnt by a more detailed analysis, particularly of the ill-defined regions.

The CASP14 assessment for T1027 was limited to residues with well-defined atomic positions across all 19 models in the NMR ensemble. In total, four regions were considered ill-defined and therefore excluded (Figure 2). This is also standard practice for many NMR protein structure validation programs, which typically only consider well-defined regions identified by the program CYRANGE^13^. ANSURR validation is different in that it requires consideration of the entire protein structure, as excluding residues will lead to nearby regions becoming artificially too flexible.

**Figure 2.**
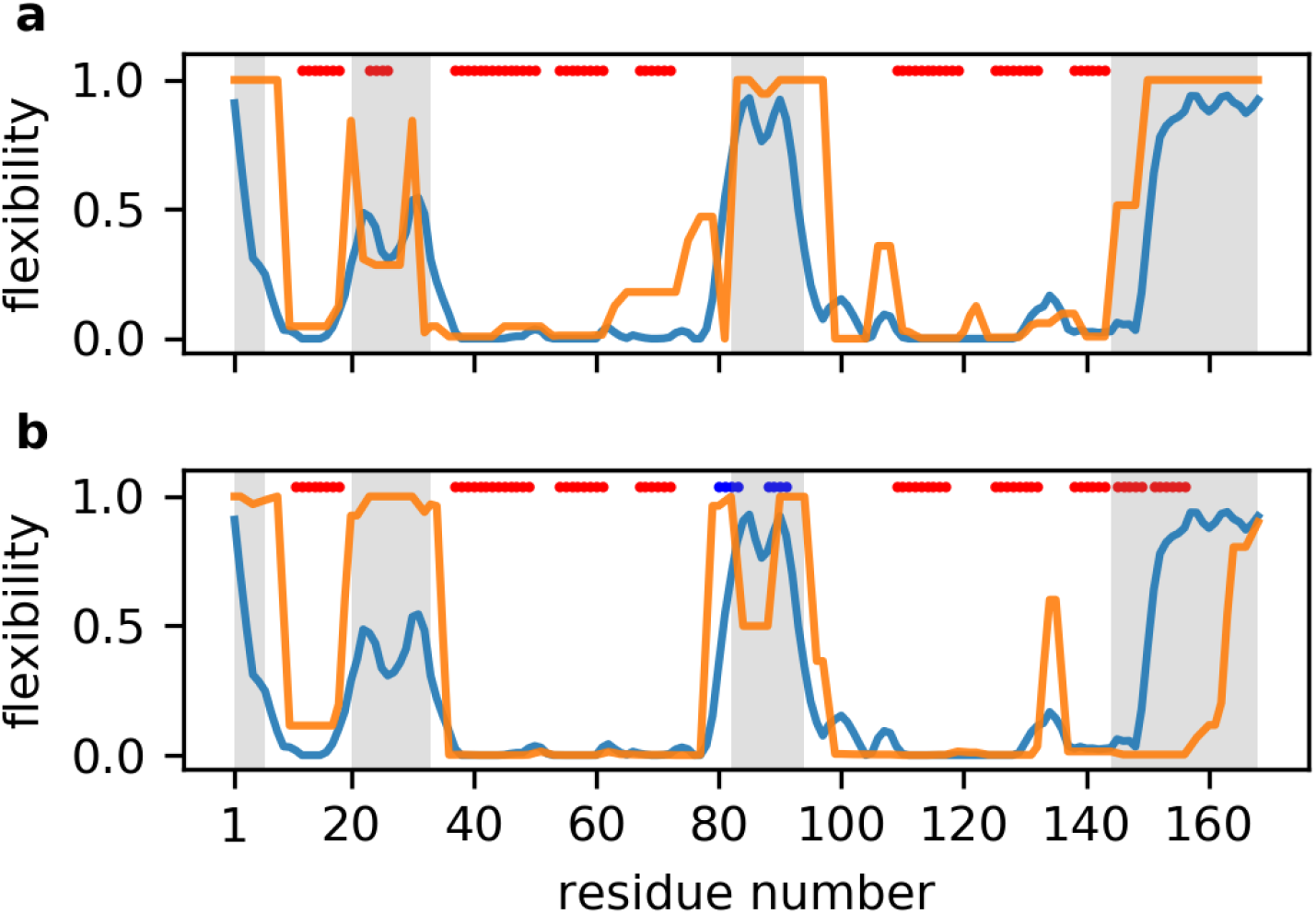
ANSURR analysis of T1027. Blue lines show the rigidity as measured by RCI based on backbone chemical shifts (BMRB 36288); orange lines show the rigidity (a) of the best scoring NMR structure (model 11 from the ensemble), and (b) of the best scoring AF2 model (model 3). Red bars at the top of each figure denote α-helical structure as assessed from the structure using DSSP, and blue bars denote β-sheet. Regions characterised as ill-defined by CYRANGE are indicated in grey.

The second ill-defined region (Figure 2, residues 20-33) is particularly interesting. The authors of the NMR structure used ^15^N relaxation dispersion and ^1^H-^15^N heteronuclear NOE data to show that this region is dynamic, and suggested it is intrinsically disordered. However, ANSURR shows it is much less flexible than the other three ill-defined regions and therefore although it is dynamic, it is not intrinsically disordered. There is also a noticeable reduction in flexibility in the center of this region. Both of these features are reflected in the computed flexibility of the NMR structure, but not in the AF2 structure. The NMR structure has a short α-helix in this region that acts to reduce the flexibility of the surrounding area, whereas this region is completely disordered in the AF2 structure (SI Figure 1a,b). Our ANSURR analysis suggests this region is flexible, in agreement with dynamic NMR measurements, but is not intrinsically disordered. ANSURR thus suggests that the helical structure is present in solution, for the majority of the time.

Chemical shifts suggest the third ill-defined region (residues 82-94) is highly disordered. There is a small reduction in flexibility between residues 86-89. This region is completely disordered in the NMR ensemble. The slight reduction in computed flexibility in this region for model 11 (shown in Fig 2a) originates from two weak hydrogen bonds, but is not observed for any of the other models from the ensemble. In contrast, the AF2 models comprise a loose β-sheet-like structure linked by a moderately rigid turn (SI Figure 1c,d). The position of the turn corresponds to the reduction in flexibility between residues 86-89 according to chemical shifts, but is more rigid. The same β-sheet-like structure is present in all five AF2 models but with variable orientation relative to the rest of the protein, perhaps indicative of dynamics. It is likely that the truth lies somewhere in between the slightly too flexible NMR structure and slightly too rigid AF2 structure. That is to say, this region in solution is dynamic and likely transitions between disorder (the NMR structure) and a loose β-sheet-like conformation (the AF2 model).

In the fourth ill-defined region (residues 144-168), the AF2 model contains an α-helix that is not present in the NMR structure. ANSURR shows this region is highly flexible and so does not support the existence of the helix. However, ^15^N relaxation dispersion and ^1^H-^15^N heteronuclear NOE data suggest this region could potentially transiently adopt secondary structure^14^. Given that chemical shifts represent a population-weighted average, it seems an α-helix in this position would not comprise the dominant conformation in solution, as suggested previously^4^.

Overall, our analysis suggests that for T1027 the experimental NMR structure is globally more accurate than the AF2 structure. However, the picture is less clear looking at the local detail. One reason for this could be that this protein is particularly dynamic and not well described by a single structure. Our ANSURR analysis also highlights the importance of validating ill-defined regions in NMR structures. Such regions can adopt a wide range of partially ordered structures.

### Target T1029

The highest scoring CASP14 prediction for T1029 had a GDT_TS of only 45, suggesting that it and all other predicted structures were highly inaccurate. However, our ANSURR analysis reveals that the target NMR structure is actually much less accurate than many of the predicted structures. In fact, 51% of the predicted structures have better ANSURR scores than the best scoring NMR model. During the preparation of this paper, it was confirmed that the NMR structure is inaccurate^4^. The NOESY peak list used to generate the original NMR structure was found to be missing many peaks present in the NOESY spectra. The NOESY peaks were carefully re-picked and used to recalculate the structure. The AF2 predictions were then used to guide refinement - referred to as “inverse structure determination” by the authors. The resulting NMR structure is very similar to the AF2 structure and has much improved ANSURR scores (green points on Fig 1). Even so, the recalculated NMR structure remained slightly less accurate than the AF2 structure. More details are presented in Supplementary Information.

### Comparison of all available human AF2 and NMR structures

Our analysis of three examples from CASP14 suggests that structures predicted by AF2 can rival or even exceed the accuracy of NMR structures. To investigate this more broadly we extended our study to compare 904 human protein structures from the recently published AlphaFold Protein Structure Database^12^ with their NMR structure counterparts from the PDB. ANSURR was used to validate each AF2 structure and each model in the corresponding NMR ensembles. To simplify the analysis of a large number of structures, correlation and RMSD scores generated by ANSURR were summed to obtain a single accuracy score, termed ANSURR score, as described previously^11^. Individual correlation and RMSD scores are provided in supplementary information.

Figure 3a shows the difference in ANSURR score between the AF2 models and the models from the corresponding NMR ensembles. AF2 structures tend to be more accurate than NMR structures, with a mean difference in ANSURR score of 28. The ANSURR score is a ranked centile score on a range from 0 to 200: this difference therefore represents a significantly better performance for AF2 compared to NMR. We have previously shown^11^ that the accuracy of the different structures within the NMR ensemble varies widely. In Figure 3b, we therefore compare the AF2 prediction to the best scoring model from the NMR ensemble. The difference in ANSURR score is now only 2, indicating a very similar overall accuracy for the two methods, though with a wide spread.

**Figure 3.**
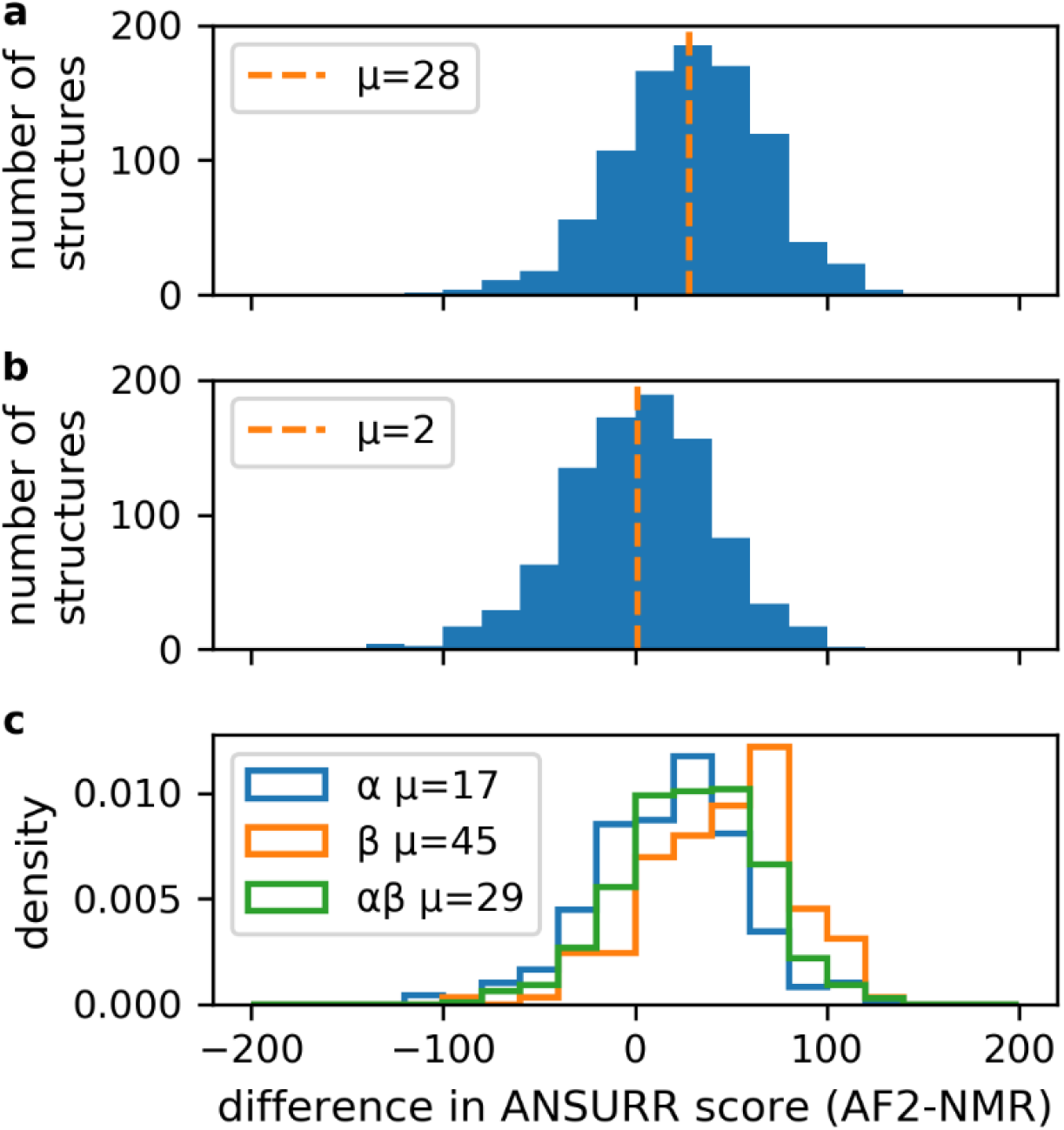
Frequency distribution for the difference in ANSURR score between the AF2 prediction and NMR structure, given as [AF2 score] – [NMR score] so that a positive difference indicates a better score for the AF2 prediction. Selection criteria are outlined in Methods. (a) Comparison of AF2 to the averaged ANSURR score for the NMR ensemble. Mean difference is 28. (b) Comparison of AF2 to the single best NMR structure in the ensemble (ie, the NMR structure with the best ANSURR score). Mean difference is 2. (c) Breakdown of the data in (a) by protein secondary structure classification as determined by DSSP, using proteins classified as α-helical, β-sheet or mixed α/β.

Fig 3c depicts the difference in ANSURR score between AF2 and NMR structures according to regular secondary structure content. We find the difference in accuracy is particularly apparent for β-sheet proteins (mean difference of 45) whereas the accuracy of α-helical proteins is closer (mean difference of 17).The difference for proteins with mixed secondary structure content falls in between (mean difference of 29). These results make sense as α-helices have limited variation in local geometry and so hydrogen bonds (important for imparting rigidity) are relatively straightforward to obtain during refinement. In contrast, β-sheets can adopt a wider range of local geometries making it more challenging to correctly resolve hydrogen bonds. We have noted this effect before^11^, finding that NMR structures often lack hydrogen bonds in β-sheets.

For a new protein target, an AF2 structure can be generated by a non-expert within a few minutes, while an NMR structure generally takes months of specialist skills and equipment. A simplistic conclusion would therefore be that AF2 is quicker, cheaper and at least as accurate, and so should be the preferred method for generation of structural models. However, the reality is more nuanced, and we approached it by looking in more detail at instances where one method represents a significant improvement over the other.

### Examples where AlphaFold structures are significantly more accurate than NMR structures

To understand why AF2 structures tend to be more accurate than NMR structures, we looked more closely at the AF2 structures that had ANSURR scores at least 50 greater than those of the NMR structures. There were 282 such structures (31% of the 904). The increased accuracy largely stemmed from AF2 models having more extensive hydrogen bond networks than NMR structures, which results in them being more rigid overall, giving them a higher ANSURR RMSD score. We have noted previously^11^ that NMR structures tend to be too floppy, and that increasing the rigidity of the NMR structure by addition of hydrogen bonds generally improves its ANSURR score. The locations of the hydrogen bonds do of course have to be correct, and AF2 provides accurate predictions of hydrogen bond locations^1^. Figure 4 provides two examples.

**Figure 4.**
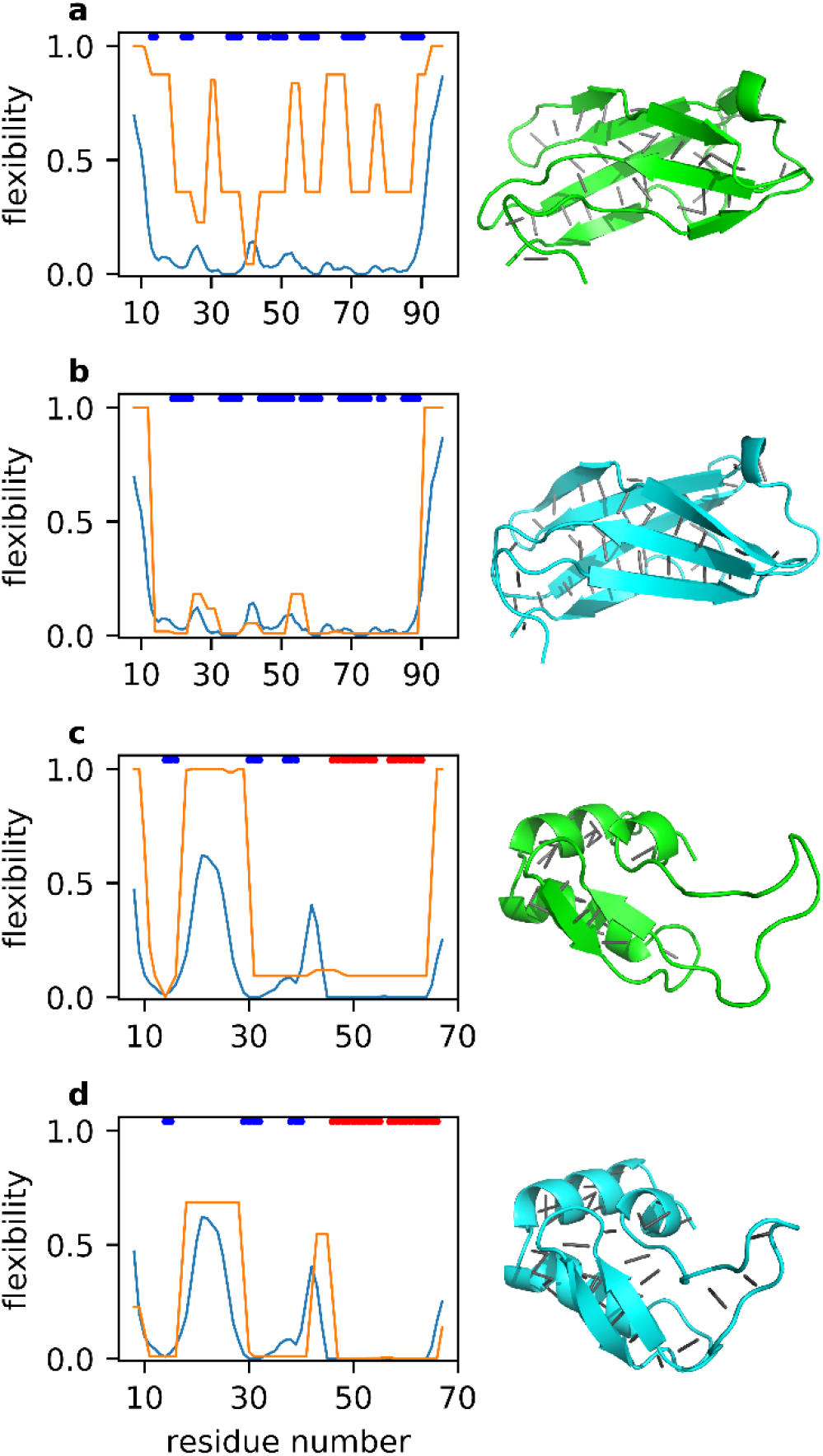
Representative ANSURR output for two proteins where the AF2 model is more accurate than the NMR structure. Each panel shows the rigidity from chemical shifts in blue, and the structure rigidity in orange. The colored bars at the top of each plot indicate regions of regular secondary structure: α-helix (red) and β-sheet (blue). The structures are shown beside each plot in cartoon representation, with backbone hydrogen bonds depicted as grey lines. (a) and (b): 20th Filamin domain from human Filamin-B. (a) is the NMR structure (PDB ID 2dlg, model 19) and (b) is the AF2 model (UniProt O75369). (c) and (d): the zinc finger BED domain of the zinc finger BED domain containing protein 1. (c) is the NMR structure (PDB ID 2ct5, model 3) and (d) is the AF2 model (UniProt O96006).

Figures 4a and 4b depict the ANSURR output for the 20th Filamin domain from human Filamin-B, a fairly rigid protein, while Figures 4c and 4d depict ANSURR output for a much more flexible zinc finger domain. For both proteins, the AF2 structure has greater rigidity, and matches better to the rigidity determined from experimental chemical shifts. For the Filamin domain (Figs 4a and b) the additional hydrogen bonds mainly define and extend the β-sheet regions better (and more correctly). The zinc finger (Figs 4c and d) has a large flexible loop between residues 16-30 which is completely lacking any backbone hydrogen bonds in the NMR structure. However, the AF2 structure contains six backbone hydrogen bonds in this region, so that the loop adopts a loose β-sheet-like conformation. These hydrogen bonds act to reduce the overall flexibility, and more specifically in a way that leads to better agreement with the flexibility obtained from chemical shifts, suggesting that they persist in solution. In summary, we suggest that the AF2 models tend to be better than NMR structures because they contain not just more hydrogen bonds but also correct hydrogen bonds that tend to persist in solution.

### Examples where NMR structures are significantly more accurate than AlphaFold structures

There were only 25 instances (3% of the 904) where NMR structures had an ANSURR score at least 50 greater than the AF2 structure. From the ANSURR output and inspection of the structures we find that there are three main reasons as to why.

First, in some cases better ANSURR scores were achieved due to differences in terminal regions that likely result from NMR measurements being performed on constructs representing only part of an entire protein e.g. a single domain. The models in the AlphaFold Protein Structure Database cover the entire sequence associated with a particular UniProt accession number whereas many NMR structures only represent some portion. As a result, terminal regions in NMR structures are likely to be more disordered/flexible than they would be as part of a larger construct, which could explain differences between NMR and AF2 structures at the C-terminal end of Figs 5a,b. An example outlining this in more detail is included in SI Figure 5. It should be noted that because we use the chemical shifts associated with an NMR structure, we are biased towards favouring NMR structures. This makes the high ANSURR scores obtained by the AF2 structures even more impressive.

**Figure 5.**
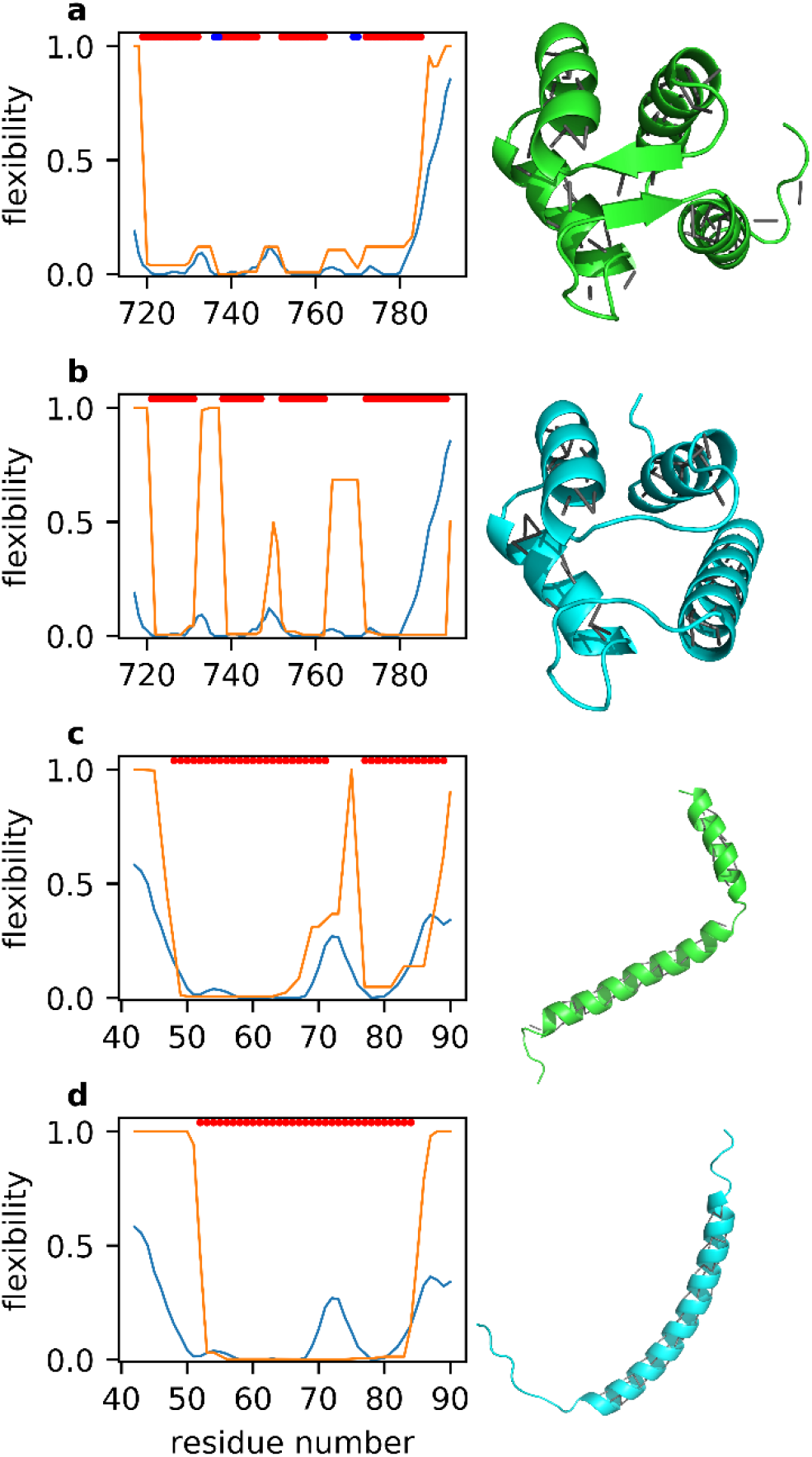
Representative ANSURR output for two proteins where the NMR structure is better than the AF2 model. Color scheme as for Figure 4. The structures are shown beside each plot in cartoon representation, with backbone hydrogen bonds depicted as grey lines. (a) and (b): EF-hand domain of human polycystin 2. (a) is the NMR structure (PDB ID 2y4q, model 3) and (b) is the AF2 structure (UniProt Q13563). (c) and (d): transmembrane and juxtamembrane domains of epidermal growth factor receptor in DPC micelles. (c) is the NMR structure (PDB ID 2n5s, model 2), and (d) is the AF2 structure (UniProt P00533),

Second, some AF2 models are missing the correct regular secondary structure. An example is shown in Figs 5a,b, where the NMR structure has a short β-sheet region which is missing in the AF2 structure. As a result, the AF2 structure is much too flexible between residues 732-738 and 763-771. We note that AF2 produces its own confidence score called per-residue local difference distance test (pLDDT). AF2 correctly indicates confidence in this particular prediction as “low” with a mean pLDDT of 66 (out of a maximum of 100, SI Figure 6a).

**Figure 6.**
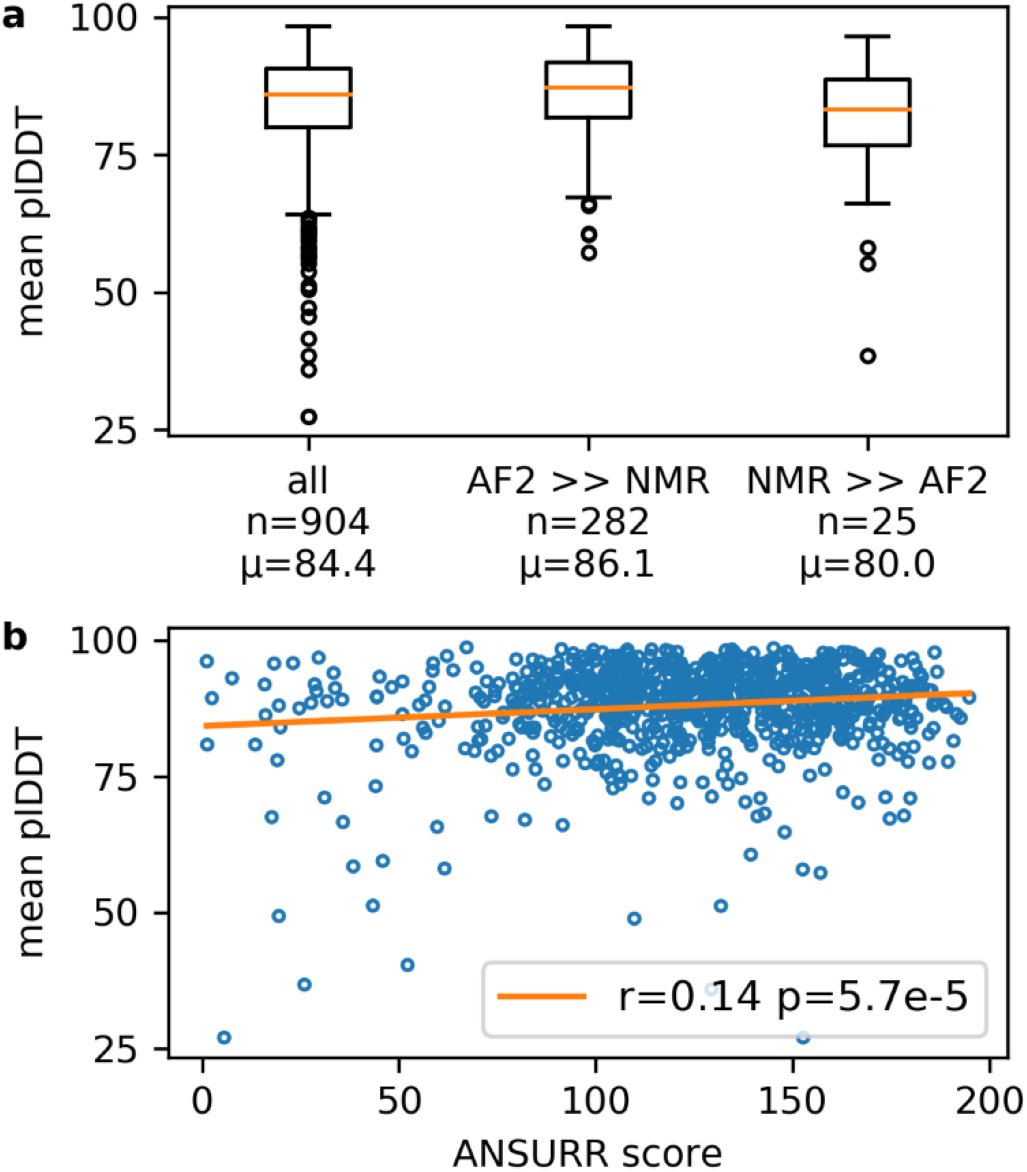
A comparison of pLDDT scores to ANSURR scores. (a) The mean pLDDT score averaged over all amino acids for each AF2 model. Statistics are shown for all AF2 models in the test set, and separately for the 282 structures in which the AF2 structure is significantly better than the NMR structure, and for the 25 structures in which the NMR structure is significantly better than the AF2 structure. The mean pLDDT score is shown below each box. (b) Correlation plot for mean pLDDT scores vs ANSURR scores for each AF2 model in the test set. The orange line is the line of best fit. Pearson’s r and the corresponding two-tailed p value are given in the legend.

Third, some AF2 models have incorrect secondary structure. Figure 5c shows the NMR structure of a membrane-associated α-helix with a break that is reflected in both the flexibility determined from chemical shifts and the computed flexibility. In contrast, the AF2 structure does not have the break, clearly in violation of the NMR data. As before, AF2 correctly indicates “low confidence” in the prediction, with a mean pLDDT of 58 and particularly low confidence in the region that should contain the break (SI Figure 6b). We speculate that AF2 will struggle to predict breaks in helices as they are less commonly observed in crystal structures (because they are difficult to crystallise or because crystallisation stabilises unbroken helices) and are therefore under-represented in the AF2 training data.

### Comparison of estimated per-residue pLDDT and ANSURR scores

Figure 6 shows two examples where the AF2 structures are less accurate than NMR structures. In both cases, AF2 had correctly identified a low confidence in the predictions, via a low mean pLDDT. We therefore carried out an analysis to see whether mean pLDDT can be used as a measure of accuracy. Figure 6a shows that the AF2 models that have significantly better ANSURR score than the NMR structures (AF2 >> NMR) have a larger mean pLDDT, whereas the AF2 models that have significantly worse ANSURR scores have a smaller mean pLDDT. However overall there is little correlation between pLDDT and ANSURR score (Fig 6b). In a paper accompanying the public release of AF2, it was demonstrated that regions with low pLDDT tend to be disordered, to the extent that pLDDT can be used as highly competitive disorder predictor^15,16^. Hence, AF2 may assign low confidence to a disordered region which ANSURR highlights as accurate because it correctly lacks structure (see Figure 4c,d as an example of how ANSURR can distinguish between regions of high flexibility and complete disorder).

## Discussion

It is already clear that the availability, simplicity and remarkable accuracy of AF2 will make it invaluable for modelling protein structures, for example for the design of drugs that work by binding to the protein. However, this is only true as long as the AF2 models are good models for the structure of the protein <u>in solution</u>. The studies presented here compare AF2 models to solution chemical shifts, and provide convincing evidence for the accuracy of AF2 models as solution structures, confirming earlier reports^17,18^. Nevertheless, there are rare occasions where the AF2 models are incorrect, likely because they do not adequately represent the dynamics of proteins in solution. Can NMR be used to identify and correct such errors?

Two reviews comparing NMR and crystal structures^6,7^ have concluded that NMR structures have the same fold as corresponding crystal structures, but are on average of lower quality. Our own analysis using ANSURR^10,11^ reached the same conclusion. An interesting point made by Andrec, et al.^7^ is that the precision of the NMR ensemble is tighter than the average distance between the NMR ensemble and the crystal structure: that is, that the most obvious measure of the “error” of the NMR structures is misleadingly small - not only are NMR structures of low quality but the error attached to them is unreliable. More recent analyses have reached similar though slightly more optimistic conclusions: thus, Schneider, et al.^19^ showed that NMR structures can be useful templates for structural models; Abaturov and Nosova^20^ showed that structural differences are minimised by collecting more NMR data; Li and Brüschweiler^21^ showed that molecular dynamics optimisation of NMR structures can make them much more comparable to crystal structures; Everett, et al.^22^ revisited the analysis of Andrec, et al. ^7^, and concluded that agreement between NMR and crystal structures is improved by using modern NMR methods; and Faraggi, et al.^23^ concluded that much of the difference may reflect genuinely increased mobility in solution. We have shown that although NMR structures are significantly too floppy by comparison to chemical shift data, crystal structures are too rigid. Indeed, numerous studies have shown that NMR structures can represent the dynamic nature of protein structures in solution better than crystal structures: for example^24,25^. These studies are of relevance to the current work, because AF2 predictions are trained on crystal structures. Thus, if NMR can be used to “correct” crystal structures to produce a more correct dynamic solution structure, it can clearly do the same also for AF2 structures.

Most AF2 structures are at least as accurate as NMR ensembles. Calculation of an AF2 prediction takes minutes and can be done with minimal training. By contrast, the calculation of an NMR structure usually takes months, and requires expensive equipment and a trained operator. It is impractical to calculate an NMR structure for every target. However, the backbone NMR assignment of small to medium sized proteins can be done almost automatically^26,27^, and permits the application of ANSURR. On the basis of the results presented here, we therefore propose that it would make sense to test the accuracy of AF2 models by carrying out semi-automated backbone assignment, followed by ANSURR. A model validated by ANSURR can be accepted as an accurate solution model (with no need for further NMR structure calculation), while models that have clear local violations need revision and would be good targets for NMR-based structure refinement of the AF2 model. Figure 7ab provides a good example of how this could be done. ANSURR shows that the AF2 model for human polycystin 2 (UniProt Q13563) is inaccurate in that it is missing a short antiparallel β-sheet present in solution. It would be straightforward to calculate a more accurate structure by starting from the AF2 model and adding additional restraints to resolve the β-sheet.

It may be argued that such a procedure biases the resulting NMR structure by imposing interatomic interactions present in the AF2 starting model. However, bias of this type is imposed on every NMR structure calculation by the use of knowledge-based restraints. The use of an AF2 model is just a more sophisticated version of a knowledge-based restraint and should be welcomed.

A complementary approach would be to produce a modified version of AF2 trained to generate more accurate solution structures, by “learning” the locations of dynamic structure. Such an approach would be enormously powerful, but would of course require the generation of appropriate training sets. The most obvious way of providing suitable training sets is via NMR chemical shifts, which carry all the information needed to characterise local dynamic regions^28,29^ and are often available from the Biological Magnetic Resonance Data Bank (BMRB)^30^. Alternatively, training data for solution structure and dynamics could be generated from molecular simulations^31^ or deep learning methods^32^.

Finally, we note that most structure calculations and structure predictions assume that the structure can be represented by a single structure. In general this seems to be true, but some of the examples discussed here suggest some element of heterogeneity, even if only in the form of folded and unfolded local structure in equilibrium. Such heterogeneity is potentially of great importance for both function and inhibition of function, and the results presented here suggest that a combination of AF2 and ANSURR would be one way to identify and characterise such equilibria.

## Methods

### A set of comparable NMR and AlphaFold structures

Each structure in the AlphaFold Protein Structure Database^12^ is indexed by a UniProt accession number. We used the Structure Integration with Function, Taxonomy and Sequence (SIFTS) resource^33^ to map the UniProt accession number of each human protein in the AlphaFold Protein Structure database to NMR structures in PDB^34^. Specifically, we used the uniprot_segments_observed.tsv SIFTS file to identify overlapping regions between the two types of structures and extracted these regions from the structure files using an in-house program. AF2 structures do not contain hydrogen atoms, so we added them using the program REDUCE v3.23^35^. We applied the following criteria to filter out NMR structures which could complicate our comparison. NMR structures needed to a) comprise only a single chain, b) have a set of backbone chemical shifts in the BMRB with at least 75% completeness, to ensure the reliability of ANSURR, and c) have at least 20 amino acid residues. The final set consisted of 904 AlphaFold/NMR structure pairs. A summary listing UniProt accession numbers and PDB IDs of the mapped AF2/NMR structures and corresponding residue ranges is provided in a supplementary text file (comparable_af2_nmr_structures.txt).

### ANSURR calculations

All ANSURR calculations were performed with ANSURR v1.1.0 (DOI 10.5281/zenodo.4984229) with the following options: re-reference chemical shifts using PANAV, include non-standard residues when computing flexibility, do not include ligands when computing flexibility. NMR structures contain multiple models (typically 20) and so we computed ANSURR scores for all models and averaged them to obtain a single ANSURR score for each PDB entry. Each AF2 structure could be mapped to multiple PDB entries. In this case we computed the average ANSURR score of the PDB entries and compared this to the average ANSURR score computed for regions taken from the AF2 structure which overlapped with the PDB entries. For example, AF2 structure O00206 was mapped to two PDB entries (5NAM and 5NAO), so we compared the average ANSURR score for the two PDB entries with the average ANSURR score for models comprising residues 623-670 and residues 623-657 from the AF2 structure. Individual ANSURR scores for all structures validated in this work are provided as supplementary text files (AF2 – af2_ansurr_scores.txt, NMR – nmr_ansurr_scores.txt). We chose not to include ligands when computing flexibility as they are not present in AF2 structures. We therefore felt that removing any ligands from NMR structures was the fairest comparison. We showed previously^11^ that ligands can cause changes in computed flexibility, but that the overall effect on ANSURR score is small: including ligands to compute flexibility for a set of 162 NMR ensembles led to a mean change in ANSURR score of only 1. Secondary structure was classified using DSSP v2.0.4^36^.

### Data availability

Source data are listed in Supplementary Information and are from publicly available databases: specifically, the Protein Data Bank (www.rcsb/org), Biological Magnetic Resonance Bank (BMRB: www.bmrb.io) and the Alphafold Protein Structure Database (https://alphafold.ebi.ac.uk). The accession codes of PDB and BMRB entries used in this study are listed in the file comparable_af2_nmr_structures. Data supporting the findings of this work are available within the paper and its Supplementary Information. The datasets generated and analysed during the current study are available from the authors upon request.

## Acknowledgements

We thank the Biotechnology and Biological Science Research Council (BBSRC) for funding to N. J. F. (BB/P020038/1).

## Author contributions

Both authors conceived the study and wrote the manuscript. N. J. F. wrote the code and did the analysis.

## References

1. Jumper, J. et al. Highly accurate protein structure prediction with AlphaFold. Nature 596, 583–589 (2021).

2. Pereira, J. et al. High-accuracy protein structure prediction in CASP14. Proteins-Structure Function and Bioinformatics 89, 1687–1699 (2021).

3. Alexander, L.T. et al. Target highlights in CASP14: Analysis of models by structure providers. Proteins-Structure Function and Bioinformatics 89, 1647–1672 (2021).

4. Huang, Y. et al. Assessment of prediction methods for protein structures determined by NMR in CASP14: Impact of AlphaFold2. Proteins Struct. Funct. Bioinf. 89, 1959–1976 (2021).

5. Williamson, M.P., Havel, T.F. & Wüthrich, K. Solution conformation of proteinase inhibitor IIA from bull seminal plasma by ^1^H nuclear magnetic resonance and distance geometry. J. Mol. Biol. 182, 295–315 (1985).

6. Billeter, M. Comparison of protein structures determined by NMR in solution and by X-ray diffraction in single crystals. Quarterly Reviews of Biophysics 25, 325–377 (1992).

7. Andrec, M. et al. A large data set comparison of protein structures determined by crystallography and NMR: Statistical test for structural differences and the effect of crystal packing. Proteins: Struct. Funct. Bioinf. 69, 449–465 (2007).

8. 8.

9. Berjanskii, M.V. & Wishart, D.S. Application of the random coil index to studying protein flexibility. Journal of Biomolecular NMR 40, 31–48 (2008).

10. Fowler, N.J., Sljoka, A. & Williamson, M.P. A method for validating the accuracy of NMR protein structures. Nature Communications 11, 6321 (2020).

11. Fowler, N.J., Sljoka, A. & Williamson, M.P. The accuracy of NMR protein structures in the Protein Data Bank. Structure 29, 1430–1439 (2021).

12. Varadi, M. et al. AlphaFold Protein Structure Database: massively expanding the structural coverage of protein-sequence space with high-accuracy models. Nucleic acids research 50, D439–D444 (2022).

13. Kirchner, D.K. & Güntert, P. Objective identification of residue ranges for the superposition of protein structures. Bmc Bioinformatics 12, 170 (2011).

14. Wu, N. et al. Solution structure of *Gaussia* Luciferase with five disulfide bonds and identification of a putative coelenterazine binding cavity by heteronuclear NMR. Scientific Reports 10, 20069 (2020).

15. Tunyasuvunakool, K. et al. Highly accurate protein structure prediction for the human proteome. Nature 596, 590–596 (2021).

16. Ruff, K.M. & Pappu, R.V. AlphaFold and implications for intrinsically disordered proteins. Journal of Molecular Biology 433, 167208 (2021).

17. Robertson, A.J., Courtney, J.M., Shen, Y., Ying, J. & Bax, A. Concordance of X-ray and AlphaFold2 models of SARS-CoV-2 main protease with residual dipolar couplings measured in solution. Journal of the American Chemical Society 143, 19306–19310 (2021).

18. Zweckstetter, M. NMR hawk-eyed view of AlphaFold2 structures. Protein Science 30, 2333–2337 (2021).

19. Schneider, M., Fu, X. & Keating, A.E. X-ray vs. NMR structures as templates for computational protein design. Proteins-Structure Function and Bioinformatics 77, 97–110 (2009).

20. Abaturov, L.V. & Nosova, N.G. Crystallographic and NMR spectroscopic protein structures: Interresidue contacts. Molecular Biology 46, 287–303 (2012).

21. Li, D.-W. & Brüschweiler, R. Protocol to make protein NMR structures amenable to stable long time scale molecular dynamics simulations. Journal of Chemical Theory and Computation 10, 1781–1787 (2014).

22. Everett, J.K. et al. A community resource of experimental data for NMR/X-ray crystal structure pairs. Protein Science 25, 30–45 (2016).

23. Faraggi, E., Dunker, A.K., Sussman, J.L. & Kloczkowski, A. Comparing NMR and X-ray protein structure: Lindemann-like parameters and NMR disorder. Journal of Biomolecular Structure & Dynamics 36, 2331–2341 (2018).

24. Tomlinson, J.H., Ullah, S., Hansen, P.E. & Williamson, M.P. Characterization of salt bridges to lysines in the protein G B1 domain. J. Am. Chem. Soc. 131, 4674–4684 (2009).

25. Ikura, M. et al. Secondary structure and side-chain ^1^H and ^13^C resonance assignments of calmodulin in solution by heteronuclear multidimensional NMR spectrocopy. Biochemistry 30, 9216–9228 (1991).

26. Würz, J.M., Kazemi, S., Schmidt, E., Bagaria, A. & Güntert, P. NMR-based automated protein structure determination. Archives of Biochemistry and Biophysics 628, 24–32 (2017).

27. Williamson, M.P. & Craven, C.J. Automated protein structure calculation from NMR data. J. Biomol. NMR 43, 131–143 (2009).

28. Dass, R., Mulder, F.A.A. & Nielsen, J.T. ODiNPred: comprehensive prediction of protein order and disorder. Scientific Reports 10, 14780 (2020).

29. Kagami, L.P. et al. b2bTools: online predictions for protein biophysical features and their conservation. Nucleic Acids Research 49, W52–W59 (2021).

30. Ulrich, E.L. et al. BioMagResBank. Nucleic Acids Res. 36, D402–D408 (2008).

31. Ramaswamy, V.K., Musson, S.C., Willcocks, C.G. & Degiacomi, M.T. Deep learning protein conformational space with convolutions and latent interpolations. Physical Review X 11, 011052 (2021).

32. Noé, F., Olsson, S., Köhler, J. & Wu, H. Boltzmann generators: Sampling equilibrium states of many-body systems with deep learning. Science 365, eaaw1147 (2019).

33. Dana, J.M. et al. SIFTS: updated Structure Integration with Function, Taxonomy and Sequences resource allows 40-fold increase in coverage of structure-based annotations for proteins. Nucleic Acids Research 47, D482–D489 (2019).

34. Burley, S.K. et al. Protein Data Bank: the single global archive for 3D macromolecular structure data. Nucleic Acids Research 47, D520–D528 (2019).

35. Word, J.M., Lovell, S.C., Richardson, J.S. & Richardson, D.C. Asparagine and glutamine: Using hydrogen atom contacts in the choice of side-chain amide orientation. Journal of Molecular Biology 285, 1735–1747 (1999).

36. Touw, W.G. et al. A series of PDB-related databanks for everyday needs. Nucleic Acids Research 43, D364–D368 (2015).

