## Supplementary Information for "The accuracy of protein structures in solution determined by AlphaFold and NMR"

### Supplementary Material

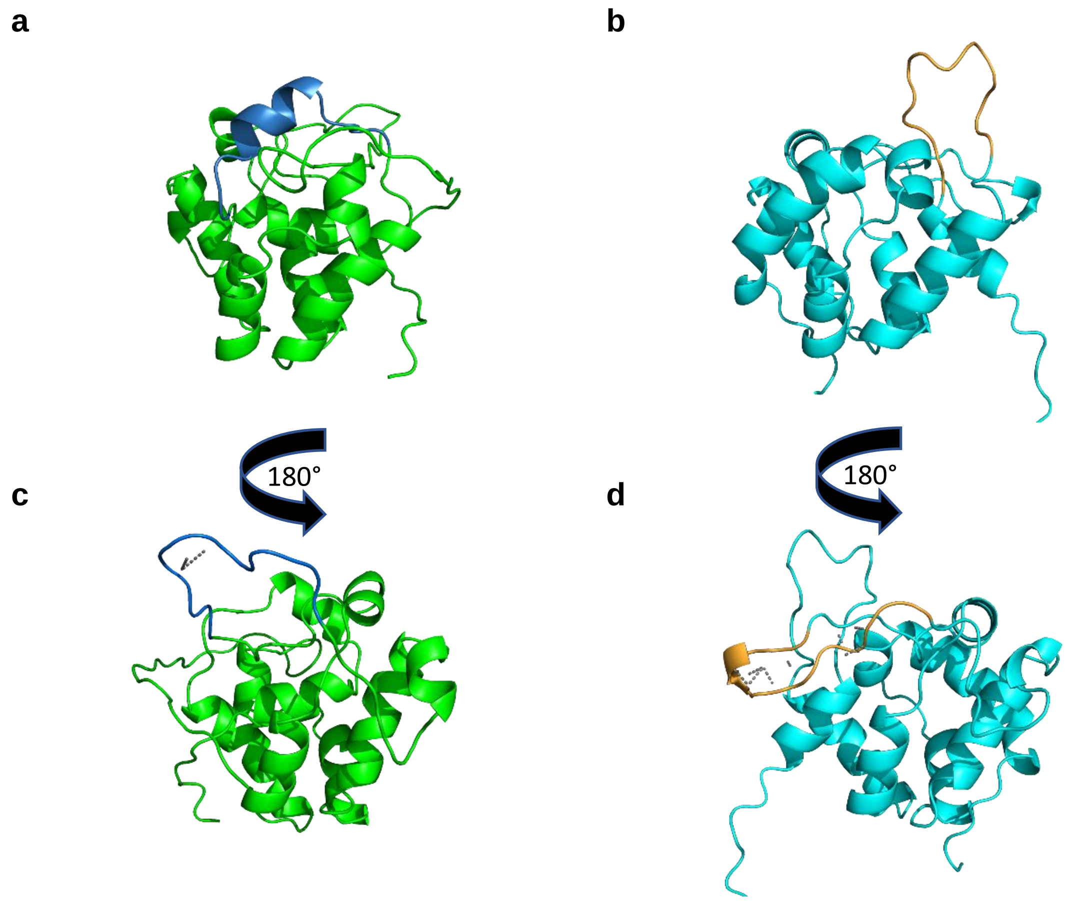


**Supplementary Figure 1.** Cartoon representations of the best scoring NMR (model 11) and AF2 (model 3) structures for target T1027 a) NMR structure with the second ill-defined region (residues 20-33) highlighted in blue. b) AF2 structure with the second ill-defined region highlighted in orange. c) NMR structure with the third ill-defined region (residues 82-94) highlighted in blue and two weak hydrogen bonds indicated by grey dashed lines. d) AF2 structure with the third ill-defined region highlighted in orange and hydrogen bonds indicated by grey dashed lines.

**ANSURR analysis for CASP14 target T1029**

ANSURR output for the original NMR structure, the AlphaFold structure and recalculated NMR structure is shown in Supplementary Figure 2. The original NMR structure contains an ill-defined region located between two $\alpha$-helices (residues 20-28). By contrast, the second helix in the AlphaFold and recalculated NMR structure extend into this region resulting in computed flexibility that is in better agreement with the flexibility determined from chemical shifts. A smaller region of ill-defined residues remains in the refined NMR structure (residues 21-25). Comparison of the refined NMR and AlphaFold structure shows significant differences in hydrogen bonding and hydrophobic contacts. These results suggest that ill-defined regions may sometimes reflect a lack of restraints rather than extensive dynamics.

Another difference between the recalculated NMR and AlphaFold structures is the region between residues 108-112. The rigidity of the AlphaFold structure in this region is imparted by a network of hydrophobic contacts formed between the sidechains of I3, L36, I42, Y104 and T110 (Supplementary Figure 3). Removing the sidechain of T110 by replacing it with an alanine results in a significant increase in computed flexibility.

All three structures appear to be too flexible between residues 46-54. In the recalculated NMR and AlphaFold structures this region is a moderately sized loop ending in a turn whereas the original NMR structure has more extensive $\beta$-sheet character (Supplementary Figure 4). Interestingly, the rigidity of the original NMR structure is a better match to the chemical shift data, although admittedly still not very good. It has already been noted that AlphaFold is less accurate when modelling loops (Feng et al., 2021). It is therefore possible that the AlphaFold and recalculated structure derived from it are inaccurate in this region. It is also possible that this region is dynamic and can convert between $\beta$-sheet-like and more extended/less structured conformations.

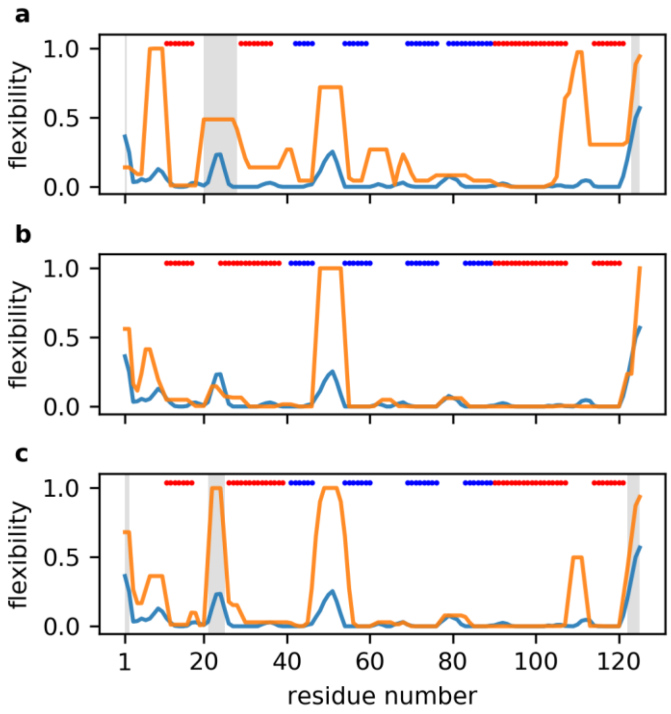


**Supplementary Figure 2.** ANSURR analysis of T1029. Blue lines show the rigidity as measured by RCI based on backbone chemical shifts (BMRB 30925); orange lines show the rigidity of (a) the best scoring model from the NMR structures used for CASP14 target T1029 (PDB 6uf2, model 10), (b) the best scoring AF2 model (model 4), and (c) the best scoring model from the NMR ensemble that was recalculated after CASP14 (PDB 7n82, model 9). Red bars at the top of each figure denote α-helical structure and blue bars denote β-sheet, as determined by DSSP. Regions characterised as ill-defined by CYRANGE are indicated in grey.

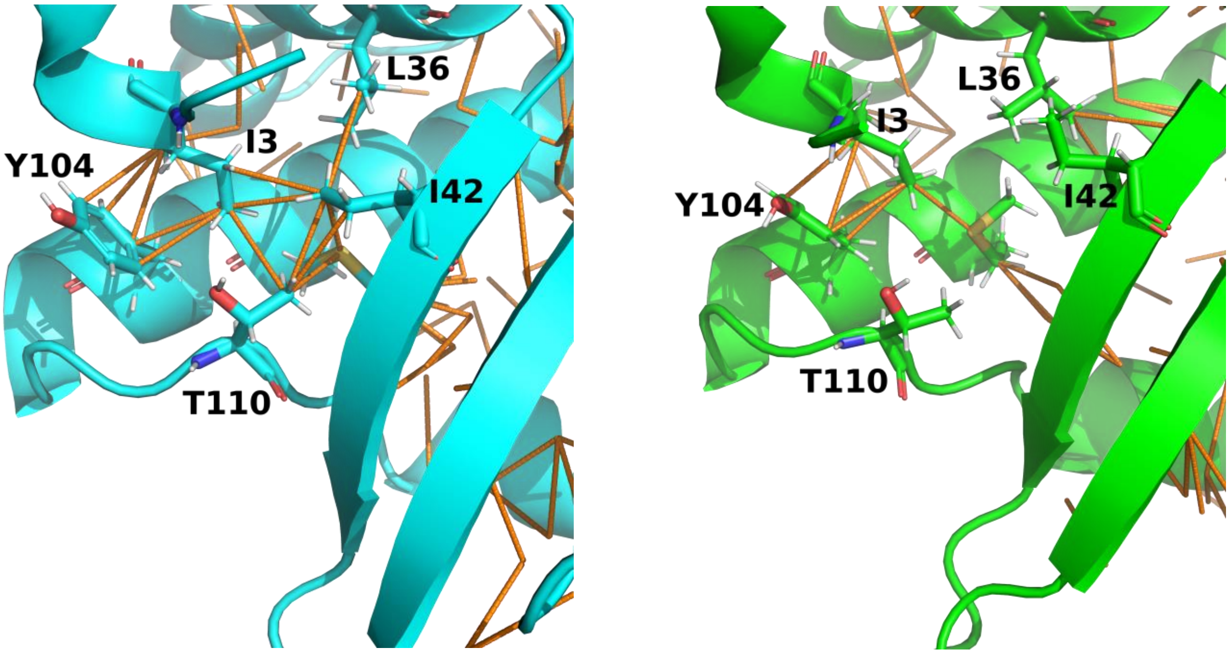


**Supplementary Figure 3.** Left: Hydrophobic contacts formed between the sidechains of I3, L36, I42, Y104 and T110 impart rigidity to residues 108-112 in the AlphaFold structure. Right: The same region in the recalculated NMR structure is missing key hydrophobic contacts, resulting in residues 108-112 being more flexible than suggested by chemical shifts.

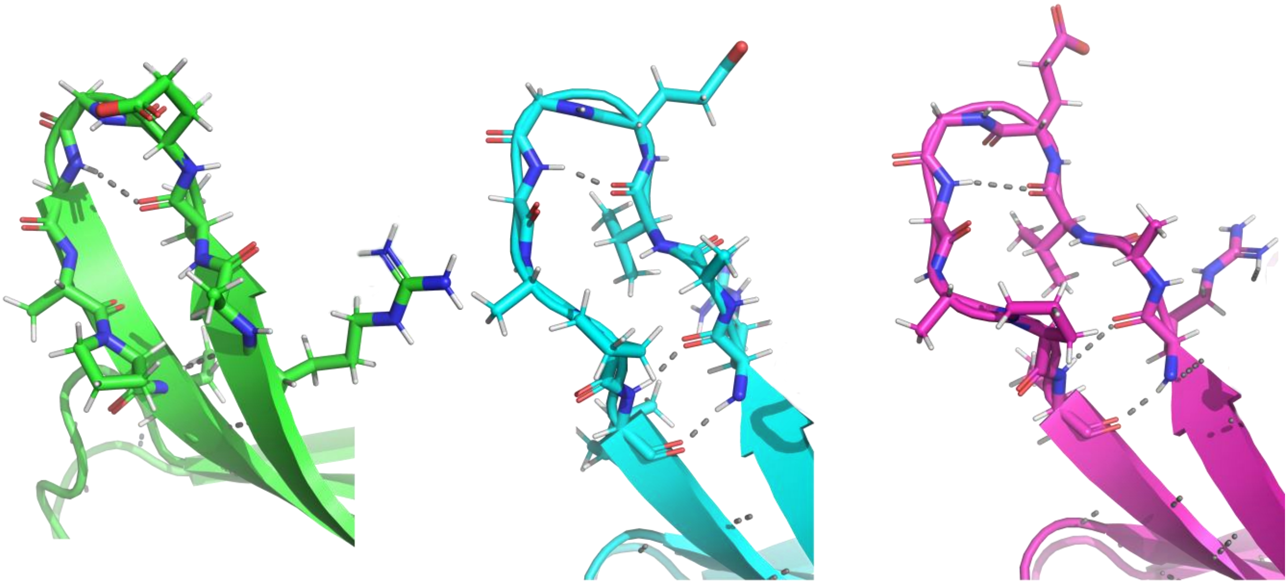

a b c

**Supplementary Figure 4.** Residues 46-54 in the a) original NMR structure, b) AlphaFold structure and c) recalculated NMR structure.

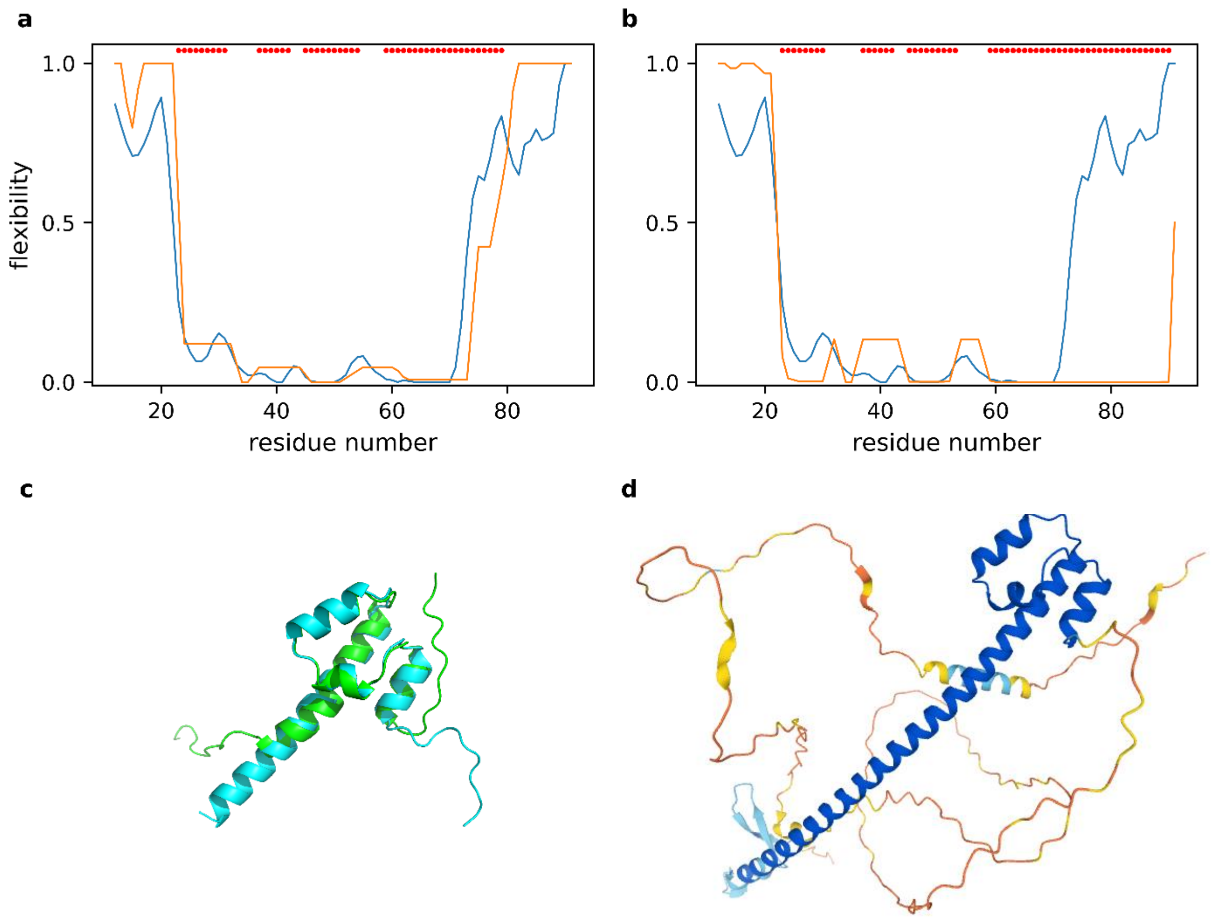


**Supplementary Figure 5.** Differences in terminal regions between NMR structures and AF2 predictions that likely result from NMR measurements being performed on constructs representing only part of an entire protein. a) ANSURR output for NMR structure of transcription factor NF-E2 subunit's DNA binding domain (PDB 2kz5, model 5). Flexibility according to chemical shifts is shown in blue, flexibility computed from the structures is shown in orange and red dots indicate α-helical regions. b) ANSURR output for AF2 prediction for the same domain (UniProt Q16621). c) cartoon representation of the NMR (green) and AF2 (cyan) structures. d) AF2 prediction for the entire protein taken from the AlphaFold Protein Structure Database.

**a** **b**

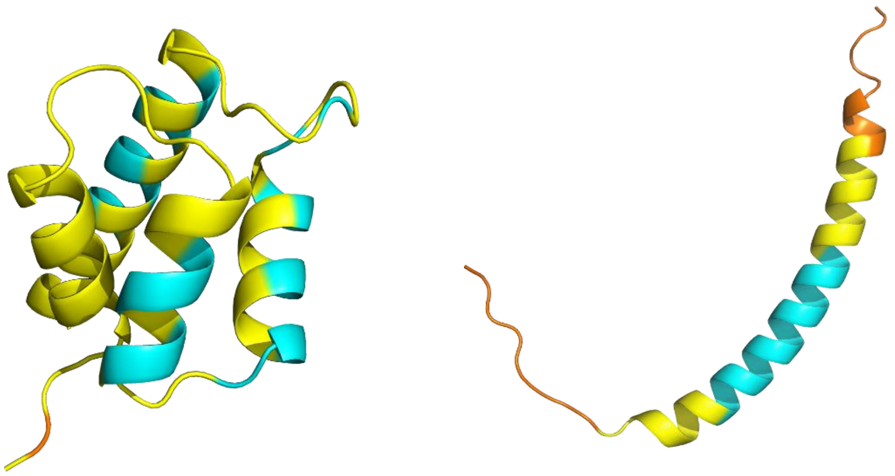

**Supplementary Figure 6**. AlphaFold predictions for a) UniProt Q13563 (mapped to PDB 2y4q) and b) UniProt P00533 (mapped to PDB 2n5s). Cartoon representation is coloured according to AlphaFold confidence metric pLDDT: "Very high" (pLDDT > 90) – dark blue, "confident" (90 > pLDDT > 70) – cyan, "low" (70 > pLDDT > 50) - yellow, "very low" (pLDDT < 50) – orange.

**Suppementary Reference**

Feng, J.-J., Chen, J.-N., Kang, W., and Wu, Y.-D. (2021). Accurate structure prediction for protein loops based on molecular dynamics simulations with RSFF2C. Journal of Chemical Theory and Computation *17*, 4614-4628.
